# Engineered pH-Sensitive Protein G / IgG Interaction

**DOI:** 10.1101/2020.12.25.424402

**Authors:** Ramesh K. Jha, Allison Yankey, Kalifa Shabazz, Leslie Naranjo, Nileena Velappan, Andrew Bradbury, Charlie E. M. Strauss

## Abstract

While natural protein-protein interactions have evolved to be induced by complex stimuli, rational design of interactions that can be switched-on-demand still remain challenging in the protein design world. Here, we demonstrate a computationally redesigned natural interface for improved binding affinity could further be mutated to adopt a pH switchable interaction. The redesigned interface of Protein G-IgG Fc domain, when incorporated with histidine and glutamic acid on Protein G (PrG-EHHE), showed a switch in binding affinity by 50-fold when pH was altered from mild acidic to mild basic. The wild type (WT) interface only showed negligible switch. The overall binding affinity at mild acidic pH for PrG-EHHE outperformed the WT PrG interaction. The new reagent PrG-EHHE will be revolutionary in IgG purification since the traditional method of using an extreme acidic pH for elution can be circumvented.

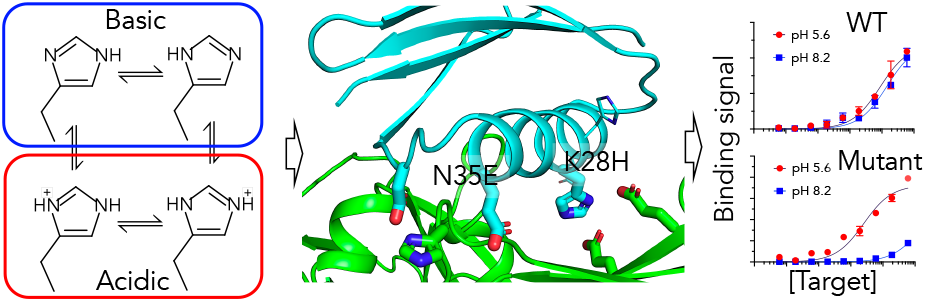

---

Purified antibodies have a wide range of applications from therapeutics to diagnostics and is a common affinity reagent in the laboratory for research. Therapeutic antibody alone has an expected market size of $125 billion by 2020.^1^ The Protein G (PrG) along with Protein A and a combination of both (Protein A/G) are commonly used in purification of antibodies. The current state-of-the-art in antibody purification is a two-step process where binding of antibody at pH ~7 followed by elution at low pH, typically 2.7. Though the acidic pH in the eluent is immediately neutralized, it is known to cause detrimental effect as the antibodies show increased structural expansion and flexibility making it prone to aggregation.^2^ Some studies have shown new structures formed at pH lower than 3.5,^3^ while others have confirmed that neutralization of elute at pH 2.7 show low recovery of functional antibody confirming certain structural changes in the antibodies are non-reversible.^4^

A key improvement in the antibody purification strategy would be to operate complete process in near physiological pH. Histidine is a special amino acid with pKa around neutral pH and has been utilized in a number of protein engineering applications for pH switchable interactions. Histidine was used at *de novo* designed interface for protein assembly to switch the oligomer in alternate conformations.^5^ The established molecular switch was as a result of protonation of histidine at mild acidic pH resulting in higher desolvation penalty and steric clashes. Similar properties at the protein-protein interaction interface resulted in 500-fold change in binding affinity between IgG and a *de novo* designed affinity reagent.^6^ Contrary to that, by utilizing the unique property of histidine, which is protonated in acidic pH and deprotonated at basic pH (Figure 1A), the protein-protein interaction can be modulated via improved charge-charge interaction in the acidic pH. Here, we exploited this idea and rationally incorporated mutations at two positions in natural PrG/IgG interaction to create a binding switch, with a tight interaction at acidic pH and significantly weaker at basic pH.

**Figure 1.**
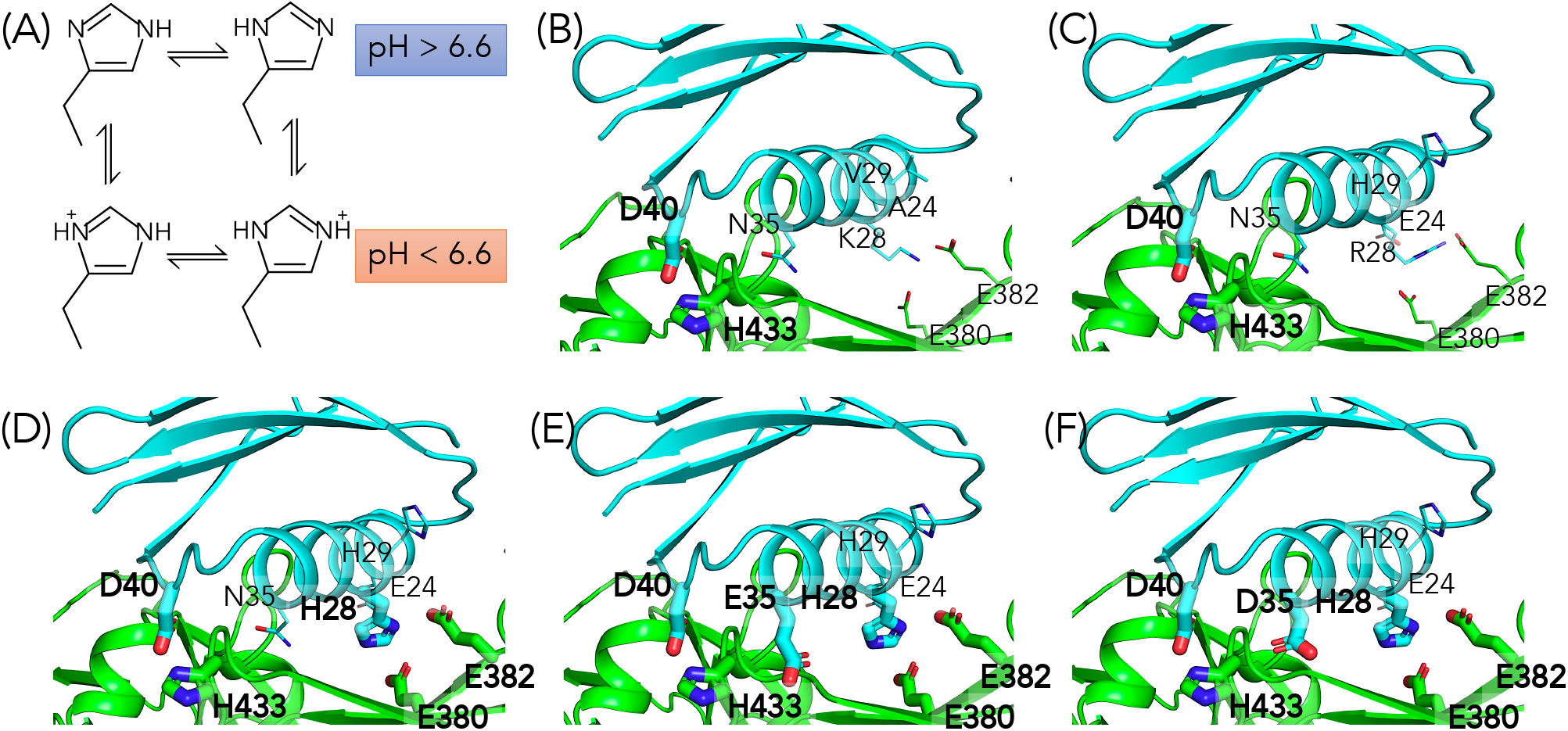
Development of a pH switchable protein-protein interaction. (A) Histidine side chain and protonated and deprotonated states at acidic and basic pH respectively. Protein G (PrG) and hIgG-Fc domain showing amino acids at the interface (B) PrG-WT, (C) PrG-ERH consisting of A24E, K28R and V29H mutations, (D) PrG-EHH consisting of K28H mutation along with A24E and V29H, (E) PrG-EHHE with added N35E to PrG-EHH and (F) PrG-EHHD with added N35D mutation to PrG-EHH. Side chains that will show perturbed interaction due to pH changes are shown in thick sticks and labeled in bold fonts.

Natural PrG (PrG-WT) binds to Fc domain of IgG with a lysine (K28) at the interface.^7^ In our earlier work, we showed K28R along with A24E and V29H mutations (the variant named PrG-ERH) showed improved expression of PrG on yeast surface and increase in binding affinity to human and rabbit IgG Fc domain.^8^ In the current work, the central arginine (R28) from the novel hydrogen bond network was chosen for mutation to histidine to give the variant PrG-EHH. Due to a shorter side chain of histidine compared to previously engineered arginine, we expected weakening of the PrG/IgG interaction. While the protonation state of arginine remains unchanged at around physiological pH (mild acidic or mild basic), we hoped that introduction of histidine at the same position will help achieve a salt bridge which is sensitive to pH changes around the pKa of ~6.6 (with a standard deviation of 1.0 depending on the environment and degree of burial of histidine side chain in consideration).^9^ Further, based on the PrG/IgG structure,^7^ we identified that the residue His-433 on the Fc domain and at the interface would also undergo change in the protonation state. Hence, position 35 on PrG, which is in close proximity to His-433, would also strongly respond to pH changes if the native asparagine (N35) is mutated to a negatively charged amino acid such as glutamate or aspartate to experience the charge-charge interaction at mild acidic pH (< pKa). Hence, the variants PrG-EHHE (with N35E) and PrG-EHHD (with N35D) were designed. The mutations were modeled using Rosetta^10,11^ and the binding energies for each complex estimated using InterfaceAnalyzer protocol^12^ (Supplementary Table S1).

Compared to the variant PrG-ERH, Rosetta predicted detrimental effect of the mutations on the binding affinity. But the overarching goal was to achieve a binding switch between mild acidic and mild basic pHs. Rosetta binding energy calculations considered neutral pH, in which the histidine residues would remain deprotonated resulting in the absence of histidine mediated charge-charge interaction at the interface. Further analysis of the structural models derived from crystal structure (PDB code 1FCC), confirmed H433 on hIgG always in close proximity to a native aspartate (D40) on PrG (Fig. 1B-F). Even though in PrG-ERH, V29H mutation was introduced, the histidine did not show stable interaction with nearest glutamate (E382) of hIgG-Fc (Figure 1C). In PrG-EHH, addition of H28 alone contributed to two more interfacial polar interactions (H28/E380 and H28/E382) that are expected to be sensitive to pH changes (Figure 1D). Further, in PrG-EHHE and PrG-EHHD, another polar interaction was created between H433 and E/D-35 (Figure 1E & 1F).

The experimental analyses of the PrG designs were carried out as described earlier.^8^ In brief, the PrG variants were displayed on the surface of yeast cells and expressed with strand 11 (S11 tag) of split GFP system.^13^ Protein expression was estimated using GFP 1-10 complementation with the S11. The binding was estimated using human IgG (hIgG) labeled with Alexafluor-647 (Jackson Immuno Research). The PrG expressing yeast cells were washed in yeast wash buffer (Phosphate buffer saline with 5 g /L bovine serum albumin), resuspended in phosphate buffer saline (PBS) and incubated overnight at 4 °C with saturating concentration of GFP 1-10 to fully complement the displayed S11 tag. The cells were washed and incubated with hIgG-APC in buffers (phosphate/citrate, phosphate or tris-HCl) at different pH (5.6, 7.0, 8.2) respectively at room temperature for approximately 1 h, washed again and resuspended in respective pH buffers. The cells were then analyzed using laser/filter combinations for GFP (excitation 488 nm, emission 530/30 nm) and APC (excitation 633 nm, emission 660/20 nm) on FACS Aria III (BD Biosciences). The population was gated for forward and side scatter (FSC-A/SSC-A) and further the gate was adjusted to include low SSC-W to eliminate any doublet.

In order to calculate the binding affinity of the PrG at different pH, serial dilutions of hIgG-APC were incubated with a fixed number of PrG expressing yeast cells (100 or 200 μL) before analyzing them using flow cytometer. The APC fluorescence response was then plotted against the concentration of hIgG and fitted with one site specific binding model to estimate the dissociation constant. This approach for calculating binding affinities has been shown to be consistent with other biochemical approaches.^14^ Initial test confirmed that PrG-WT and PrG-ERH shows negligible pH effect even though there is a histidine mediated (H433) polar interaction with aspartate (D40) on PrG. When the central arginine (R28) was mutated to histidine, the variant showed a very distinct pH effect and a large shift in binding interaction (Supplementary Figure 1). N35E or N35D that can experience polar interaction with H433, showed further enhancement of the pH-mediated binding switch when changed from mild acidic to mild basic or (neutral pH) (Supplementary Figure S1). The estimation of binding affinity at pH 5.6 and pH 8.2 showed <2-fold switch in PrG-WT while PrG-EHHE and PrG-EHHD showed >50 and >20-fold respectively (Figure 2A-C, Table 1). Based on the dose-response plot, PrG-EHHE showed >80% of maximum binding at acidic pH and ~10% at pH 8.2 at an IgG concentration of 200 nM (Figure 2D-E). The new variants PrG-EHHE and PrG-EHHD also showed improved display on yeast cells confirming enhanced expression compared to PrG-WT (Figure 2F).

**Figure 2.**
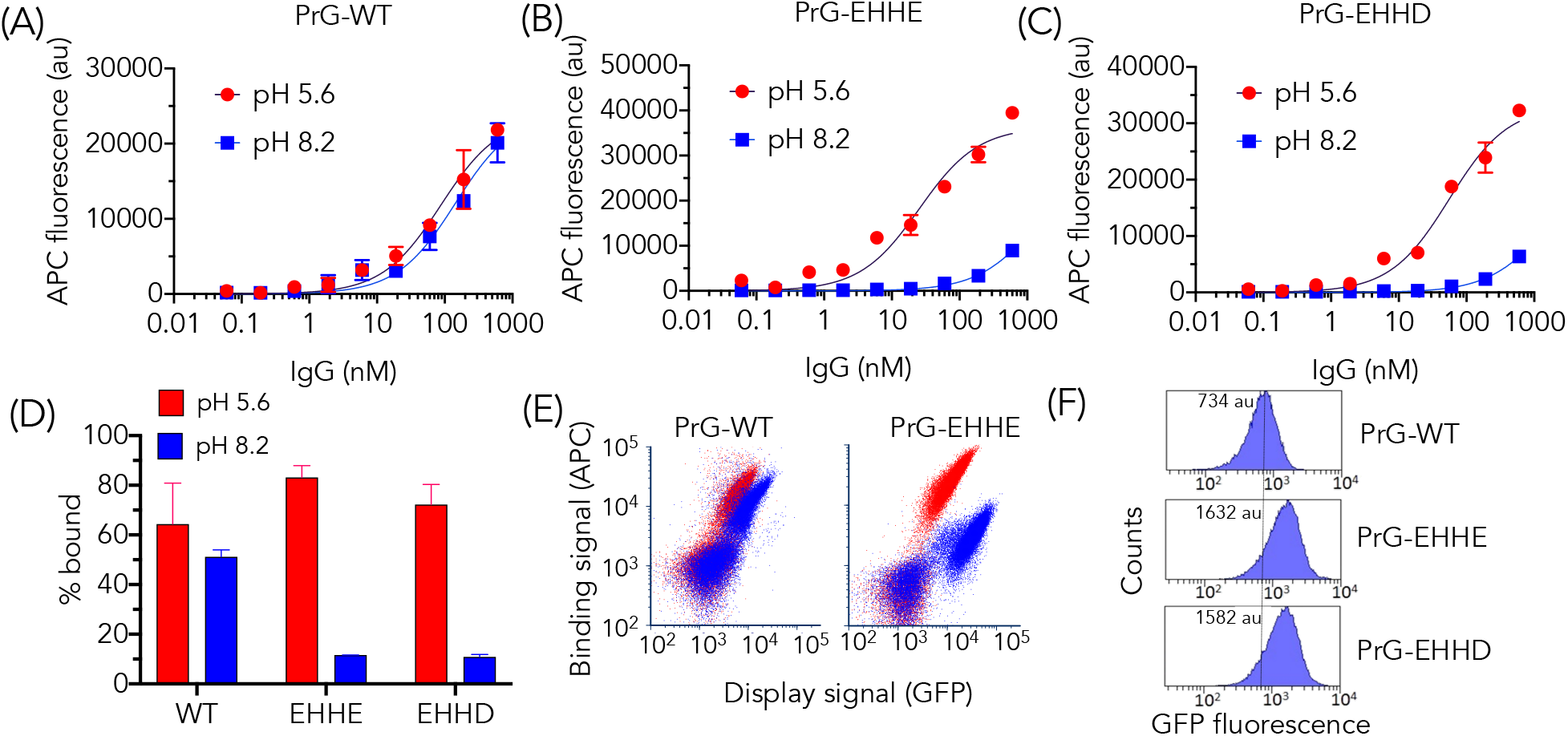
Analyzing effects of structure-based designed mutations on PrG-IgG interactions. (A) PrG-WT, (B) PrG-EHHE and (C) PrG-EHHD variants and dose-response with human IgG (hlg G) at acidic and basic pH. The plots represent mean and standard deviation from two independent experiments. One-site specific binding model was used for the curve fitting. (D) Percent hIgG bound (at 200 nlMt input concentration) to PrG variants at acidic and basic pH. (E) Scatter plot of yeast cells displaying PrG variants, PrG-WT and PrG-EHHE, and showing binding with hIgG at acidic pH (red dots) and basic pH (blue dots). (F) Display level of PrG variants on yeast. The number in the box refers to mean fluorescence level (arbitrary fluorescence units or au) of the yeast population displaying PrG.

**Table 1.** One-site binding analysis and estimation of dissociation constant (Kd) at acidic and basic pHs for PrG-IgG interaction. R^2^ value for the curve fitting is shown in parentheses.

|  | pH 5.6<br>Kd (nM) | pH 8.2<br>Kd (nM) | Fold<br>change |
| --- | --- | --- | --- |
| PrG-WT | 88 (0.96) | 145 (0.96) | <2 |
| PrG-EHHE | 25 (0.95) | 1344 (0.99) | 53 |
| PrG-EHHD | 56 (0.98) | 1471 (0.98) | 26 |

Overall, the new PrG variants show superior display, binding (at acidic pH) and binding switch upon changing the pH from mild acidic to mild basic. In conclusion, the histidine-mediated interaction is dependent on the buffer pH. Our step-by-step approach to mutate residues at the PrG/IgG interface showed major improvement in the binding switch after addition of only two mutations. While the purification of IgGs have remained challenging due to inevitable inclusion of very low acidic pH step, these reagents which operate around the physiological pH can give a new alternative and improvement in the antibody purification schemes.

## ASSOCIATED CONTENT

### Supporting Information

Table S1: Rosetta binding energy calculation for PrG variants and human IgG-fc domain

Figure S1: Measurement of binding affinity for PrG variants and human IgG at acidic, neutral and basic pH.

## ACKNOWLEDGMENT

The work was authored under Triad National Security, LLC (“Triad”) Contract No. 89233218CNA000001 with the U.S. Department of Energy. This research used computational resources provided by the Los Alamos National Laboratory (LANL) Institutional Computing Program (under w19_proteng to R.K.J.), which is supported by the U.S. Department of Energy National Nuclear Security Administration under Contract No. 89233218CNA000001. GFP S1-10 domain was a kind gift from Geoffrey Waldo (LANL).

## ABBREVIATIONS

IgG: Immunoglobulin G;
PrG: Protein G;
GFP: Green Fluorescent Protein;

## Supplementary Information

**Supplementary Table S1.** Rosetta binding energy calculation of PrG and human IgG. Mean of Top 3 binding energy and standard deviation.

| Variant | Binding energy (REU) |
| --- | --- |
| PrG-WT | -47.8±0.5 |
| PrG-ERH | -52.9±0.6 |
| PrG-EHH | -50.7±0.2 |
| PrG-EHHE | -48.4±0.4 |
| PrG-EHHD | -48.3±1.0 |

**Supplementary Figure S1.**
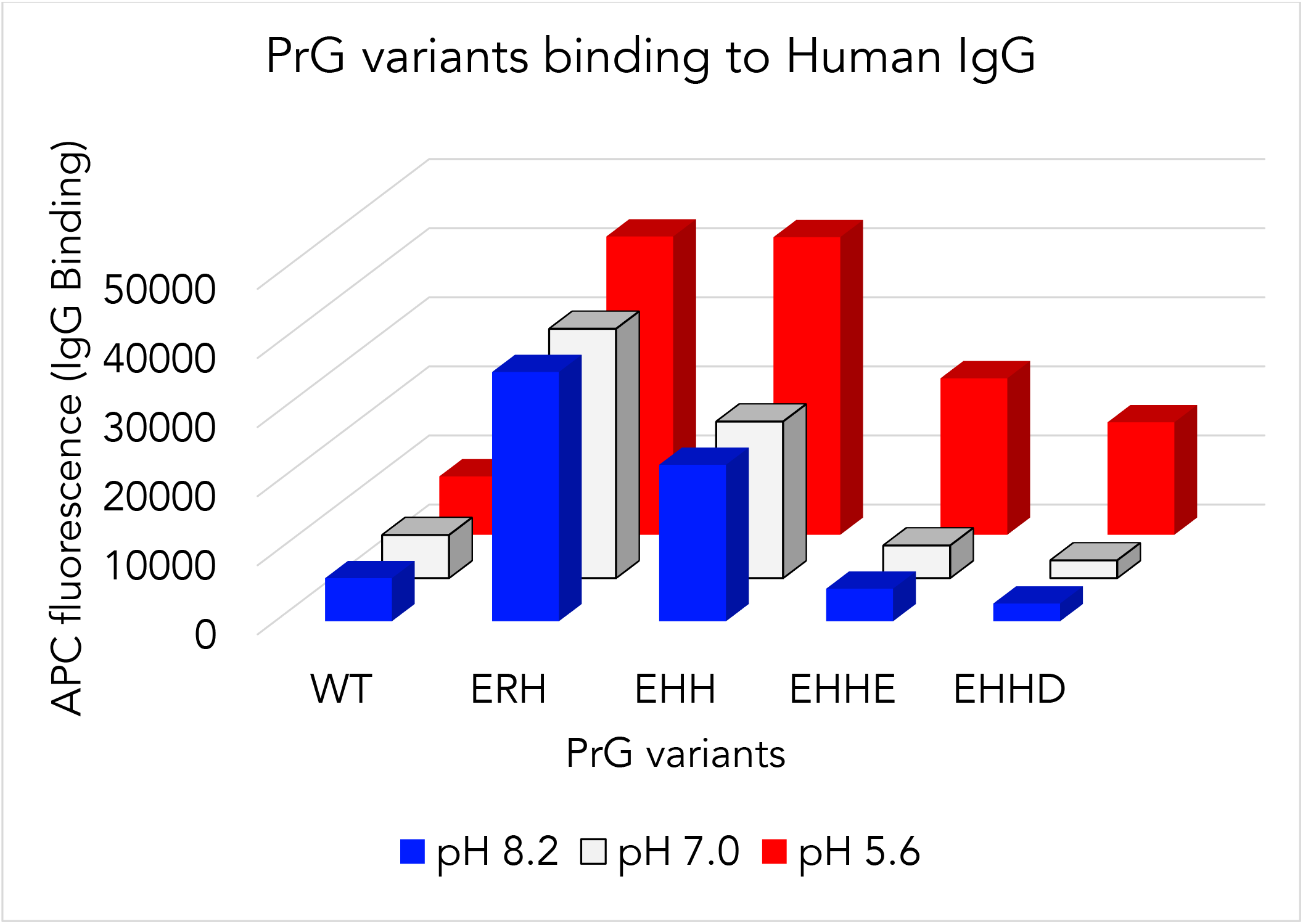
pH dependent binding of PrG variants and human IgG. APC fluorescence was indicative of IgG binding and the experiment was performed at 600 nM of IgG and PrG variants were displayed on yeast cells.

